# Socioeconomic Disparities and Sexual Dimorphism in Neurotoxic Effects of Ambient Fine Particles on Youth IQ: A Longitudinal Analysis

**DOI:** 10.1101/207589

**Authors:** Pan Wang, Catherine Tuvblad, Diana Younan, Meredith Franklin, Fred Lurmann, Jun Wu, Laura A Baker, Jiu-Chiuan Chen

## Abstract

Mounting evidence indicates that early-life exposure to particulate air pollutants pose threats to children’s cognitive development, but studies about the neurotoxic effects associated with exposures during adolescence remain unclear. We examined whether exposure to ambient fine particles (PM_2.5_) at residential locations affects intelligence quotient (IQ) during pre-/early-adolescence (ages 9-11) and emerging adulthood (ages 18-20) in a demographically-diverse population (N = 1,360) residing in Southern California. Increased ambient PM_2.5_ levels were associated with decreased IQ scores. This association was more evident for Performance IQ (PIQ), but less for Verbal IQ, assessed by the Wechsler Abbreviated Scale of Intelligence. For each inter-quartile (7.73 μg/m^3^) increase in one-year PM_2.5_ preceding each assessment, the average PIQ score decreased by 3.08 points (95% confidence interval = [−6.04, −0.12]) accounting for within-family/within-individual correlations, demographic characteristics, family socioeconomic status (SES), parents’ cognitive abilities, neighborhood characteristics, and other spatial confounders. The adverse effect was 150% greater in low SES families and 89% stronger in males, compared to their counterparts. Better understanding of the social disparities and sexual dimorphism in the adverse PM_2.5_-IQ effects may help elucidate the underlying mechanisms and shed light on prevention strategies.

## Introduction

Intelligence is a broad collection of cognitive abilities including reasoning, problem solving, attention, memory, knowledge, planning, and creativity sub-served by different parts of the brain. Intelligence quotient (IQ), a global measure of intellectual development, is an important determinant of national wealth and economic growth (1). It is estimated that a single point change of IQ could bring a gain of $55 billion to $65 billion (in year 2000 dollars) for a single birth cohort of US population (2). At the individual level, childhood IQ is a powerful predictor of later-life socioeconomic success (3). Although the brain size has reached 90% of adult size by age 5 (4), development of efficient brain structure and networks in early childhood continues into adolescence. There is an increasing recognition that IQ can change significantly during adolescence (5).

Adolescence, defined by the World Health Organization (6) as the period from ages 10 to 19 (after childhood and before adulthood), is a transition stage characterized by many significant biological and social changes. Human growth during adolescence is greatly influenced by changes in hormone production and neuroendocrine response (7) with the beginning of reproductive lifespan, while the developing brain is undergoing further remolding of gray matter (e.g., cortical thinning) (8) and white matter (e.g., continuing myelination of axons) (9). The growing adolescents start to disengage from their parents and exert more autonomous control on their own decisions and actions. These biological and social changes not only suggest that plasticity in IQ development continues with interactions among brain, behavior, and social context, but that adolescent brains are also vulnerable to environmental insults from various neurotoxins. As the brain network matures by the end of adolescence (4, 10), IQ is expected to remain relatively stable until the advent of aging during late adulthood.

Environment in general can explain up to 50% of individual difference in IQ, with its resulting influence depending on socioeconomic context (11) and age (12). Research on environment-mediated IQ effect is thus important as such knowledge may help identify potentially modifiable factors and develop timely intervention to reduce disparities in cognitive development. While there has been extensive research on IQ development and social adversities in the family and school environments (13–15), influences of physical environments are understudied.

Exposure to ambient particulate air pollutants, including PM_2.5_ (particulate matter [PM] with aerodynamic diameter <2.5 μm), has emerged as a novel environmental neurotoxin affecting brain development in children (16). The hypothesized link of child intellectual development with early-life PM exposures has been examined in several birth cohorts (17–27), including four based in the US and three from Poland, China, and Italy. Although most of the reported findings generally showed a negative association between PM exposure and IQ in children, each of these birth cohort studies included only one-time assessment on intellectual development. One small longitudinal study (28) compared children living in highly-polluted Mexico City (n=20) and the control group (n=10) from a clean-air area (matched on age and socioeconomic status), and reported in their post-hoc analyses the difference in IQ at baseline disappeared after one year of follow-up when the matched cohort became 8 years old. Therefore, it remains unclear whether PM exposure could still exert adverse effect on intellectual development during adolescence. The primary aim of our current study was to examine the adverse effect of PM_2.5_ on IQ, using longitudinal data spanning a 12-year period. Because previous studies have been underpowered to assess the potential heterogeneity in the reported associations, our secondary aim was to evaluate whether the putative neurotoxic adverse effect on intellectual development during adolescence, if any, could vary by sex and family socioeconomic status (SES) based on a relatively large sample (N=1360).

## Materials and Methods

### Participants

Participants were drawn from the University of Southern California (USC) Risk Factors for Antisocial Behavior (RFAB) twin study. RFAB is a prospective longitudinal study of the interplay of genetic, environmental, social, and biological factors on the development of antisocial behavior from pre-adolescence to early adulthood. Participating families were recruited from communities in Los Angeles and surrounding counties, with the resulting sample representative of the socio-economically-diverse multi-ethnic population residing in the greater Los Angeles area (29). To date, five waves of data have been collected from 780 twin pairs (N=1,569 in total including triplets). Study protocols were approved by the USC Institutional Review Board. Informed consents were obtained from all participants (after reaching adulthood) or their parents/guardians (during pre-adolescence).

The current study utilized IQ data collected from the RFAB cohort during pre-/early-adolescence (aged 9-11) and emerging adulthood (aged 18-20). Our analytic sample was limited to participants with at least one valid IQ score and a corresponding estimate of air pollution exposure, plus complete data on major sociodemographic characteristics (including age, gender, race/ethnicity and family SES). A total of 1,360 subjects (from 687 families) were retained in the main analyses, including 810 tested during pre-/early- adolescence only, 170 during emerging adulthood only, and 380 at both age periods. These three groups did not differ in the distributions by sex, race/ethnicity, or family SES (S1 Table). Subjects tested with higher IQ scores at baseline were more likely to participate in the follow-up, but their IQ scores were no different from those only tested during the emerging adulthood. The PM_2.5_ exposure 1-year before the baseline testing was slightly lower among subjects tested twice, as compared to those not participating in the second testing (20.28 ± 2.82 vs. 20.59 ± 2.53; *p* = .06), but there was no statistically significant difference in the PM_2.5_ exposure estimate at the follow-up between the two groups assessed during emerging adulthood.

### Assessment of IQ

IQ was measured using the Wechsler Abbreviated Scale of Intelligence (WASI) (30). The WASI provides a quick and reliable assessment of an individual's verbal, nonverbal, and general cognitive functioning. The WASI yields two standardized scores: Verbal IQ and Performance IQ. Verbal IQ (VIQ) is based on subtests Vocabulary and Similarities, whereas Performance IQ (PIQ) is based on subtests Block Design and Matrices. Correlations between PIQ and VIQ ranged from 0.48 (pre-/early- adolescence) to 0.56 (emerging adulthood) in the current study. The six-month test-retest reliability (n = 60) was satisfactory for both PIQ (*r* = 0. 79) and VIQ (*r* = 0.78).

### Estimation of Particulate Matter Exposure

*Residential Location Data and Geocoding*. Residential addresses for RFAB families were prospectively collected through self-reports every 2 to 3 years. Addresses were geocoded using services of the USC Spatial Sciences Institute, which successfully matched residences by exact parcel locations or specific street segments for 98.6% of participating families. The remaining addresses were checked for correctness using Google Earth and thereafter geocoded.

*Spatiotemporal modeling for PM*_*2.5*_. Daily PM_2.5_ (PM with aerodynamic diameters < 2.5μm) concentrations were obtained from U.S. EPA Technology Transfer Network for the years 2000 to 2014. A spatiotemporal model based on the measured PM_2.5_ concentrations was constructed (with 10-fold cross-validation R^2^=0.74-0.79) to estimate monthly average PM_2.5_ concentrations for each subject’s geocoded residential location (see the A2 in S1 File for more details). A time series of monthly PM_2.5_ concentrations for the 2000-2014 period was constructed and monthly estimates were aggregated to represent PM_2.5_ exposure 1-, 2-, and 3-years preceding each IQ assessment.

### Relevant Covariates

To control for potential confounding, four groups of covariates were examined: (A) age, gender, race/ethnicity, family SES, and parents’ cognitive abilities; (B) parent-reported neighborhood quality, neighborhood SES (nSES), traffic density and neighborhood greenspace; (C) CALINE4-estimated total annual nitrogen oxides (NO_x_) and temperature/humidity; (D) parent-level risk factors (operationalized as maternal smoking during pregnancy and parental perceived stress). Covariates (A) and (B) were considered as the most relevant confounders as they were known to predict IQ and also likely influence where people chose to reside (and thus their exposure to ambient PM_2.5_). More details about the selection and measurement of covariates are available in A3 of S1 File.

### Statistical Analyses

Three-level mixed-effects models regressing IQ scores (Full-Scale IQ, VIQ and PIQ separately) at each assessment on the corresponding PM_2.5_ exposures and accounting for both within-family (random intercepts and slopes of PM_2.5_ effects by families) and within-individual (random intercepts by individual) covariance were constructed as the base models. These models were then adjusted for two sets of covariates incrementally: (1) individual and family characteristics—age (as a continuous variable or dichotomized as pre-/early- adolescence vs. emerging adulthood), sex, race/ethnicity, family SES, and parental cognitive abilities; and (2) neighborhood characteristics—nSES, neighborhood greenspace (1000m radius buffer, 1-year preceding test), traffic density (300m radius buffer), and parent-reported neighborhood quality. We conducted further sensitivity analyses by adding the following covariates to the fully adjusted models: ambient temperature and humidity (1-year preceding); total annual NO_x_; and parental risk factors.

Three separate pre-planned moderation analyses were conducted to examine whether the putative PM_2.5_ effects on IQ varied by age (pre-/early- adolescence vs. emerging adulthood), sex, and SES levels (continuous), based on the interaction term between exposure and the putative moderator, each entering the fully adjusted model one by one. All the analyses were implemented using SAS 9.4.

## Results

### Descriptive Statistics

Participants’ IQ scores were on average 101.62 (VIQ, SD = 17.93) and 100.25 (PIQ, SD = 17.98) during pre-/early- adolescence (9.59 ± 0.58 years); 104.47 (VIQ, SD = 16.01) and 102.71 (PIQ, SD = 16.01) during emerging adulthood (19.44 ± 1.07 years). About 99% of participants during pre-/early- adolescence and 78% during emerging adulthood were exposed to PM_2 5_ (1-year preceding the IQ assessment) levels exceeding the EPA annual standard (12ug/m^3^).

Population characteristics by quartiles of PM_2.5_ (Table 1) and Full-Scale IQ (Table 2) at the study baseline (i.e., the first valid IQ assessment) were examined. The decrease of quartiles of PM_2.5_ exposure across age reflected the higher ambient PM_2.5_ levels in earlier years of testing. Compared to their counterparts, those with relatively higher PM_2.5_ exposures were mostly Hispanics and Blacks, from lower quality neighborhoods (characterized by lower nSES, lower greenness, more negative perception of neighborhood quality and higher annual NOx), residing in locations with higher temperature and relative humidity, and children whose parents reported maternal smoking during pregnancy, displayed poorer cognitive abilities, and perceived more stress. On the other hand, children with lower IQ score at baseline were more likely to be Hispanics, Black, and mixed racial/ethnicities; grow up in lower SES households; have parents perceiving more stress, smoking during pregnancy and demonstrating lower cognitive abilities; and reside in locations with lower neighborhood qualities and higher relative humidity. For population characteristics by quartiles of VIQ and PIQ, please refer to S2 and S3 Tables.

### Main-effect of PM_2.5_ on IQ Scores

In the base models, higher one-year average PM_2.5_ predicted lower scores in the full-scale IQ, VIQ, and PIQ (Table 3). Although PM_2.5_ exposures were still negatively associated with full-scale IQ and VIQ in the adjusted analyses, none of these associations reached statistical significance. However, the observed adverse PM_2.5_ effects on PIQ were evident in the adjusted models. For each inter-quartile (7.73 μg/m^3^) increase in 1-year PM_25_, the average PIQ score decreased by 3.08 points (95% CI = [−6.04, −0.12]) in the mixed-effect model accounting for within-family/within-individual correlations, demographic characteristics, family SES, parents’ cognitive abilities, perceived neighborhood quality, nSES, traffic density, and measure of greenspace (Adjusted Model-II). The observed adverse PM_2.5_-PIQ effect remained robust in sensitivity analyses with further statistical adjustment for temperature and humidity (Sensitivity Model-1), total annual NO_x_ (Sensitivity Model-II), and parental stress and maternal smoking during pregnancy (Sensitivity Model-III).

Additional analyses on 2- and 3-year average PM_25_ exposure effects on IQ (full-scale; VIQ; PIQ) revealed a fairly similar pattern of associations across different temporal scales of exposure (S1 Fig). Post-hoc analyses were also conducted to explore the possibility of differential impact of PM_2.5_ on each component score of PIQ (Block Design; Matrix Reasoning)or VIQ (Vocabulary; Similarities). We found the negative PM_2.5_-PIQ association primarily reflected the adverse effect on Matrix Reasoning. Interestingly, although the negative PM2.5-VIQ associations were not statistically significant (S1 Fig), we found evidence for adverse effects on VIQ Similarities present for both 1-y (*p* = .04) and 2-year (*p* = .02) PM_25_ exposures (S1 Fig). Annual NO_x_ exposure also predicted lower IQ scores in the crude analyses (S4 Table), but their associations were largely abolished in the adjusted analyses (S4 Table).

**Table 1.**
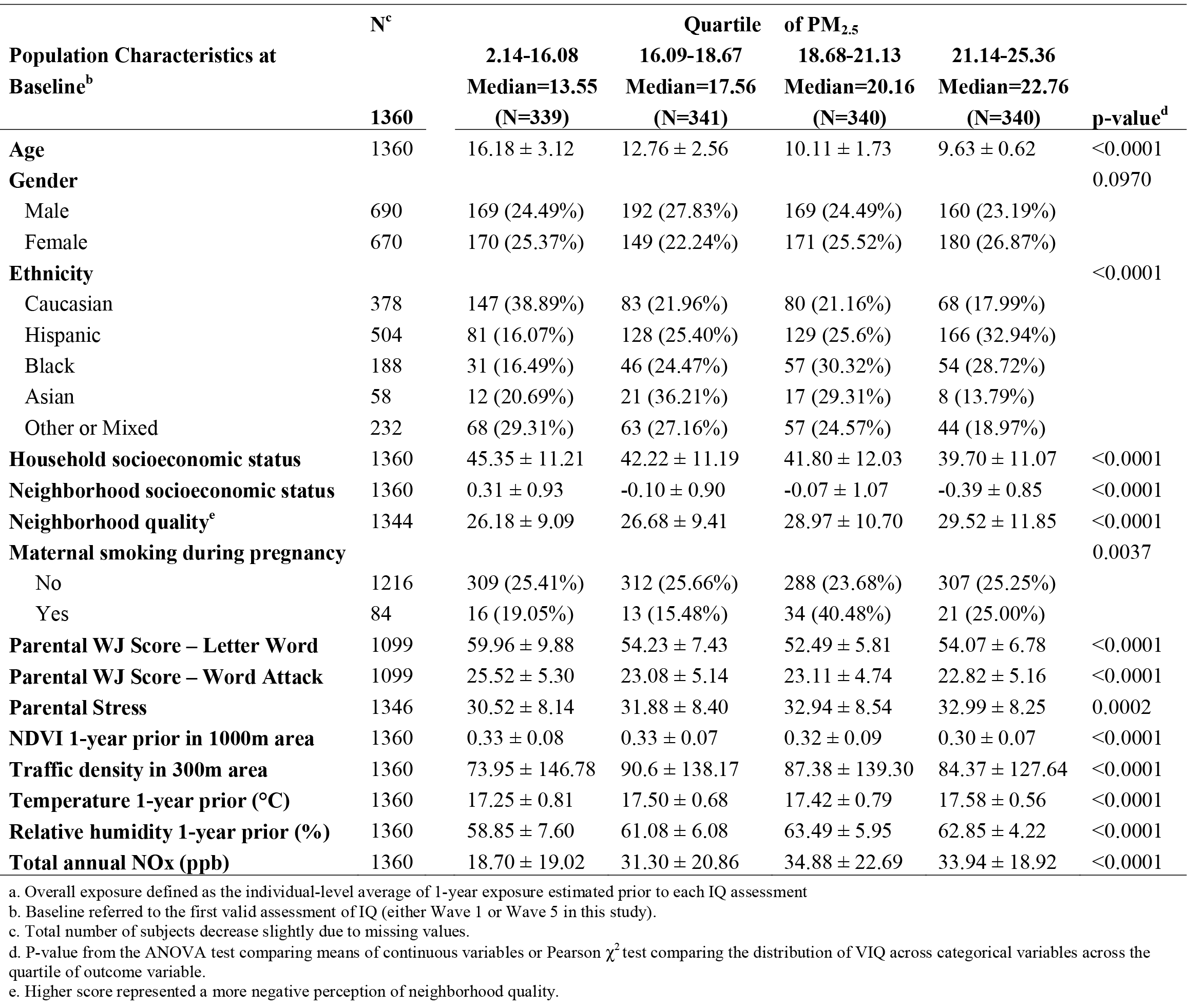
Population Characteristics in Relation to the Overall^a^ PM_2.5_ Exposure 1-Year Prior to IQ Assessment

**Table 2.** Population Characteristics at Baseline in Relation to Full-Scale IQ

| Population Characteristics | N<sup>a</sup> | Quartile of IQ |  |  |  | p-value<sup>b</sup> |
| --- | --- | --- | --- | --- | --- | --- |
|  |  | 45-92<br>Median=83<br>(N=351) | 93-103<br>Median=99<br>(N=327) | 104-114<br>Median=109<br>(N=345) | 115-149<br>Median=121<br>(N=337) |  |
| **Age** | 1360 | 10.73 ± 3.01 | 10.55 ± 2.85 | 10.83 ± 3.27 | 10.91 ± 3.40 | 0.4707 |
| **Gender** |  |  |  |  |  | 0.5498 |
| Male | 690 | 176 (25.51%) | 164 (23.77%) | 168 (24.35%) | 182 (26.38%) |  |
| Female | 670 | 175 (26.12%) | 163 (24.33%) | 177 (26.42%) | 155 (23.13%) |  |
| **Ethnicity** |  |  |  |  |  | <0.0001 |
| Caucasian | 378 | 28 (7.41%) | 58 (15.34%) | 106 (28.04%) | 186 (49.21%) |  |
| Hispanic | 504 | 182 (36.11%) | 156 (30.95%) | 110 (21.83%) | 56 (11.11%) |  |
| Black | 188 | 73 (38.83%) | 49 (26.06%) | 40 (21.28%) | 26 (13.83%) |  |
| Asian | 58 | 11 (18.97%) | 16 (27.59%) | 20 (34.48%) | 11 (18.97%) |  |
| Other or Mixed | 232 | 57 (24.57%) | 48 (20.69%) | 69 (29.74%) | 58 (25.00 %) |  |
| **Household socioeconomic status** | 1360 | 36.65 ± 9.70 | 39.14 ± 11.31 | 44.43 ± 10.73 | 48.94 ± 10.36 | <0.0001 |
| **Neighborhood socioeconomic status** | 1360 | -0.54 ± 0.59 | -0.16 ± 0.90 | 0.07 ± 0.91 | 0.39 ± 1.17 | <0.0001 |
| **Neighborhood quality<sup>c</sup>** | 1344 | 30.01 ± 12.02 | 27.58 ± 10.67 | 27.4 ± 9.39 | 26.35 ± 8.96 | <0.0001 |
| **Maternal smoking during pregnancy** |  |  |  |  |  | <0.0001 |
| No | 1216 | 296 (24.34%) | 295 (24.26%) | 308 (25.33%) | 317 (26.07%) |  |
| Yes | 84 | 34 (40.48%) | 21 (25.00%) | 21 (25.00%) | 8 (9.52%) |  |
| **Parental WJ Score – Letter Word** | 1099 | 53.34 ± 8.23 | 54.21 ± 8.06 | 55.51 ± 7.68 | 57.33 ± 7.67 | <0.0001 |
| **Parental WJ Score – Word Attack** | 1099 | 22.31 ± 6.30 | 22.75 ± 5.09 | 24.02 ± 4.57 | 25.43 ± 3.81 | <0.0001 |
| **Parental Stress** | 1346 | 33.70 ± 8.57 | 32.60 ± 8.05 | 31.95 ± 8.72 | 30.07 ± 7.76 | <0.0001 |
| **NDVI 1-year prior in 1000m area** | 1360 | 0.29 ± 0.06 | 0.31 ± 0.08 | 0.33 ± 0.08 | 0.35 ± 0.09 | <0.0001 |
| **Traffic density in 300m area** | 1360 | 90.85 ± 146.9 | 88.42 ± 148.94 | 86.99 ± 142.76 | 69.87 ± 109.65 | 0.1801 |
| **Temperature 1-year prior (°C)** | 1360 | 17.41 ± 0.70 | 17.47 ± 0.72 | 17.46 ± 0.77 | 17.41 ± 0.72 | 0.6110 |
| **Relative humidity 1-year prior (%)** | 1360 | 62.62 ± 5.92 | 61.64 ± 6.27 | 61.23 ± 6.65 | 60.76 ± 6.37 | 0.0010 |
| **Total annual NOx (ppb)** | 1360 | 32.80 ± 21.83 | 31.25 ± 22.38 | 28.74 ± 21.58 | 26.01 ± 19.21 | 0.0002 |

a. Total number of subjects decrease slightly due to missing values.
b. P-value from the ANOVA test comparing means of continuous variables or Pearson $\chi^2$ test comparing the distribution of VIQ across categorical variables across the quartile of outcome variable.
c. Higher score represented a more negative perception of neighborhood quality.

**Table 3.** Associations between PM_25_ and IQ Measures

| Models | N<sup>a</sup> | Full-Scale IQ<br>$\beta$ (95% CI)<sup>b</sup> | VIQ<br>$\beta$ (95% CI)<sup>b</sup> | PIQ<br>$\beta$ (95% CI)<sup>b</sup> |
| --- | --- | --- | --- | --- |
| Crude Analysis | 1360 | -2.46 (-3.48, -1.44)* | -1.66 (-2.76, -0.56)* | -2.14 (-3.16, -1.12)* |
| Adjusted Model I<sup>c</sup> | 1093 | -1.93 (-4.75, 0.89) | -1.37 (-4.39, 1.65) | -2.91 (-5.83, 0.01) |
| Adjusted Model II<sup>d</sup> | 1085 | -2.00 (-4.84, 0.84) | -1.42 (-4.48, 1.64) | -3.08 (-6.04, -0.12)* |
| **Sensitivity Analyses** |  |  |  |  |
| Sensitivity Model I<sup>e</sup> | 1085 | -1.84 (-4.86, 1.18) | -1.14 (-4.37, 2.09) | -3.50 (-6.62, -0.38)* |
| Sensitivity Model II<sup>f</sup> | 1085 | -2.08 (-4.96, 0.80) | -1.76 (-4.84, 1.32) | -3.01 (-5.99, -0.03)* |
| Sensitivity Model III<sup>g</sup> | 1042 | -2.05 (-4.87, 0.77) | -1.13 (-4.17, 1.91) | -3.66 (-6.62, -0.70)* |

\* $P < .05$
a. Total number of participants differed because of missing values.
b. Estimate reflected the change in each IQ score and the resulting 95% confidence interval per each inter-quartile range (IQR) increase in PM<sub>2.5</sub>.
c. Adjusted for age, gender, ethnicity, family SES and parents' cognitive abilities.
d. Adjusted Model I + neighborhood SES, self-reported neighborhood quality, traffic density (300m) and neighborhood greenness (1000m, 1-year preceding).
e. Adjusted Model II + temperature and relative humidity 1-year prior to test.
f. Adjusted Model II + total annual NO<sub>x</sub>.
g. Adjusted Model II + parental stress and maternal smoking during pregnancy.

### Moderation Roles of Socio-Demographic Characteristics

Results of our moderation analyses showed that the adverse PM_2.5_ effects on PIQ were not uniform across socio-demographic characteristics (upper panel, Fig 1). Sex and family SES both significantly modified the association between PM_2.5_ and PIQ score (interaction *p* < .01 for both moderators), with exposure conferring 150% stronger influence in males (β = −4.68, 95% CI = [−7.90, −1.47]) than in females (β = −1.87, 95% CI = [−4.89, 1.16]); and 89% stronger in low SES families (β = −3.83, 95% CI = [−6.98, −0.69]) than in high SES families (β = −2.03, 95% CI = [−6.12, 2.36]). Although the adverse PM_2.5_-PIQ effect (β = −3.27; 95% CI = [−6.44, −0.10]) at age 9-11 was 74% greater than the corresponding estimate (β = −1.88; 95% CI = [−6.12, 2.36]) during emerging adulthood, this observed difference by age did not reach statistical significance (interaction *p* = .49),

The moderation analyses of VIQ did not reveal remarkable findings, except for a statistically significant interaction (*p* = .03) between gender and PM_2.5_ (lower panel, Fig 1). Our results suggested that the PM_2.5_-VIQ effect might be qualitatively different between males (β = − 2.16; 95% CI = [−5.5, 1.18]) and females (β = 0.78, 95% CI = [−2.37, 3.93]), albeit an overlap between these two CIs (please refer to Knezevik (31) for an explanation of why a significant difference could have overlapping CIs).

**Fig 1.** Plot of regression coefficients and 95% confidence intervals for the association between PM_2.5_ 1-year prior to test and the IQ scores, moderation by age, sex, and family socioeconomic status (RFAB Cohort 2000-2014). The gray reference band in each IQ subscale represented the 95% CI of the final-adjusted main effect of PM_2.5_ on that IQ score. Significant moderation was highlighted in yellow.

## Discussion

To our knowledge, this is the first longitudinal study examining the effects of ambient air pollutants on IQ spanning two different developmental stages pre-/early-adolescence (aged 9-11) and emerging adulthood (aged 18-20). We found strong evidence for a decreased PIQ score with higher exposure to ambient PM_2.5_ estimated at residential locations, even after adjusting for socio-demographic factors, spatial characteristics of residential neighborhoods, and parents’ cognitive abilities. The corresponding associations with VIQ were less evident. The adverse PM_2.5_-PIQ effect was much greater in low SES families and in males, indicative of socioeconomic disparities and sexual dimorphism in the developmental neurotoxicity of PM_2.5_ exposure.

The observation of stronger adverse PM_2.5_ effects on IQ among RFAB participants growing up in low SES families offers a useful view-scope to unify the findings reported in the extant literature (11 studies from 7 birth cohorts with individual-level exposure data) on PM-IQ associations (A4 in S1 File). For those 4 studies conducted outside the US (19, 20, 23, 25), differences in PM characterization and primary exposure source may explain the discrepancies in reported associations. Of the 7 US-based studies, 6 reported a statistically significant association between early-life exposure to PM and low performance of IQ testing in children. These included 4 studies based in the Columbia Center for Children’s Environmental Health Birth Cohort, which included children of minority (Black or Dominican-American) women primarily with low SES (74% families with annual family income <$20,000) and residing in a community where traffic and residential heating were major exposure sources (18, 21, 24, 26). The other 3 studies, despite all having been based in the greater Boston area and employing the same approaches to estimating residential exposure at birth locations, yielded very different results. In the Project Viva (22), neither black carbon nor PM_2.5_ exposure predicted lower IQ in children (with an average age of 8) of relatively well-off (73% with annual family income >$70,000) and well-educated parents (68% maternal/ 63% paternal education ≥college). For the other two studies including mothers primarily of minorities and/or with limited educational attainment (69-82% with maternal education≤ high school), PM_25_ was associated with low full-scale IQ in boys of school age (6.5 ± 0.98 years) (27), whereas black carbon exposure predicted low Matrices score on the Kaufman Brief Intelligence Test at 8-11 years of age (17). All these study findings point to the importance of population social context (32) for designing epidemiological studies and interpreting data on developmental neurotoxicity of ambient air pollutants.

Our finding of socioeconomic disparities in the adverse PM_2.5_-PIQ effect has important implications for future research on the environmental neurosciences in neurodevelopmental toxicity of particulate air pollutants. First, PM_2.5_ exposure and socioeconomic adversities may have converged on common pathways with resulting exacerbated neurotoxicity, although the exact models for their respective mechanistic actions remain unclear. Possible brain regions and structures with shared vulnerability may include hippocampus (33, 34), prefrontal cortex (35, 36), and cerebral white matter (24, 37). Second, high-SES families may provide their children with more exposure to advantageous experiences (e.g., early-life educational resources), which could partly off-set the brain damage from PM_2.5_ exposure. Third, although our analyses accounted for parental cognitive abilities, low-SES families may not be able to engage in activities with parental nurturance critical for cognitive development. Fourth, growing up in low SES families indicates the possibility of concurrent exposures to other psychosocial and environmental stressors (e.g., violence exposure, early onset of alcohol use) adversely affecting IQ development. Better understanding of the causes of socioeconomic disparities in PM neurotoxicity will not only shed light on the mechanistic pathways, but also help identify more susceptible populations who can benefit the greatest from environmental regulation, social policies (e.g., reducing family poverty; early education program), or family interventions (e.g., parental caring behaviors).

Although PIQ and VIQ were moderately correlated, the adverse PM_2.5_-IQ effect was statistically significant for PIQ only (primarily affecting the Matrix Reasoning component). This divergence may reflect a more detrimental impact of PM on fluid cognitive abilities. Fluid intelligence (Gf) refers to the capabilities to reason and solve novel problems, in contrast to crystallized intelligence (Gc), another factor of intelligence concerning acquired knowledge, skills and experiences (38, 39). This classical distinction laid the theoretical foundation for the development of PIQ and VIQ. It is interesting to note that our *ad hoc* analyses (S1 Fig) also showed that increased PM_25_ (1- and 2-year average) exposure was associated with decreased scores in the VIQ subtest Similarity, a measure intended for Gc but actually tapping into Gf (likely more than the PIQ subtest Block Design, a spatial visualization task) as it relies upon the ability to abstract common patterns beyond the knowledge of words and their meanings (40). Because Gf is more reliant on and sensitive to lesions to frontal lobe than Gc (41–45), the differential PM_2.5_ effect on fluid intelligence implies possible damage to frontal brain networks, which was supported by the emerging data from neurotoxicological and neuroimaging studies. For instance, persistent glial activation in frontal cortex was demonstrated in mouse models with early-life exposure to concentrated ambient ultrafine particles (46). *In utero* exposure to a low concentration of diesel exhaust also altered the neurochemical monoamine metabolism in prefrontal cortex (47). In a birth cohort study based in Rotterdam, the Netherlands, early-life exposure to PM_2.5_ was associated with cortical thinning in the frontal lobe at age 9 (48).

Two recent studies have reported adverse PM effects on IQ (27) and working memory (49) assessed in school age were stronger in boys than girls, although none of the exposure interaction with sex was statistically significant. Our study showed that the adverse PM_2.5_ effects on both PIQ and VIQ scores assessed during early adolescence and emerging adulthood were stronger in males than females (interaction p-value < .05; Fig 1), despite female RFAB participants being more likely to reside in locations with higher PM_25_ (3^rd^ and 4^th^ quartiles in Table 1). Multiple biological differences may help explain the observed differences between males and females in observed adverse PM_2.5_-IQ effects in the current study. Neurotoxicologists have documented sexually dimorphic neurobehavioral responses to various environmental chemicals (e.g., dioxin, bisphenol-A), a phenomenon often inferred as an indicator for exposure-induced endocrine-disrupting effects on the brain, largely through interference with the actions of gonadal hormones (50). Animal studies support the neuroendocrine disruption with inhaled exposure to particles (51, 52), but the mechanisms underlying sexual dimorphism in neurotoxicity may also involve neurobiological pathways with exposure interacting with sex-linked genes (53). Although earlier studies did not show clear evidence for sex differences in general intelligence (54), new findings support the presence of cognitive sex differences depending on task characteristics and contextual experience (55). However, studies relating pubertal sex hormones to cognitive abilities in adolescents have yielded mixed results (56, 57). Nonetheless, our findings give strong rationale for future studies to investigate whether sexual dimorphism is also present in other neurodevelopmental and behavioral effects of ambient air pollutants. Better understanding of the neurobiological processes underlying the sexual dimorphism in the PM_2.5_-IQ effect may inform better sex-sensitive intervention strategies to reduce harmful environmental exposures to optimize the brain-behavioral health for both men and women.

Our moderation analyses revealed no statistical interaction of exposure effect by age group, despite the fact that the adverse PM_2.5_-PIQ effect was 74% stronger in pre-/early-adolescence than in emerging adulthood. Behavior genetic research has reported that environmental contribution to IQ variation decreases across age (12, 58). As neural structure and network approach maturation by the end of adolescence (4, 10), IQ of young adults may be less subject to environmental influences. Previous studies have shown that the use of neurotoxic agents, such as alcohol and other drugs, posed more threats to memory and memory-related brain function in adolescents than adults (59). However, given a relatively small sample (n=510) assessed during emerging adulthood, our results must be viewed with caution, as they did not necessarily mean that the neurotoxic threats of ambient air pollutants disappeared once into adulthood. Hippocampal damage with cognitive impairments was previously documented in mice with long-term inhaled exposure to concentrated PM_2.5_ starting in youth (33). Future studies with larger samples could help clarify this important uncertainty in the adverse PM_2.5_-IQ effect during the transition into young adults.

The strengths of our study included its base in Southern California with wide exposure contrast, sampled from a population with rich diversity in race/ethnicity, sex and family SES, and the inclusion of repeated IQ assessment for longitudinal analyses. This unique sample and prospective longitudinal design provided adequate power to investigate heterogeneity in the PM-IQ associations across age, sex, and SES. Nonetheless, there are several limitations that should be considered. First, we caution the interpretation of selective PM_2.5_-PIQ effect. Because our assessment of IQ was based on the WASI (an abbreviated Wechsler intelligence scale, rather than the full scale), some significant domains (e.g., working memory; processing speed) presumably sensitive to PM_2.5_ neurotoxicity were not captured in our analyses. Second, although we were able to conduct longitudinal analyses, the inference of our results was based on the statistical assumption of data missing at random given the unbalanced data structure with repeated measures. Third, we were not able to study prenatal exposure effects, because extensive monitoring of PM_2.5_ data were not available until after 1999, while the birth years of the cohort ranged from 1990-1995. The relative contribution to adverse PM_2.5_-IQ effects by exposure in early life versus adolescence needs to be investigated further. Fourth, our analyses only included the estimate of PM_2.5_ mass, and we did not study the specific neurotoxicity of PM_25_ constituents (e.g., metals; organic chemicals). Fifth, while PM_2.5_ estimates based on spatiotemporal interpolation of monitored concentrations were statistically cross-validated, there are expected non-differential measurement errors in such estimates, which would likely have attenuated the observed associations.

In this first longitudinal study with repeated cognitive assessment, we found lower PIQ scores in youth living in locations with higher exposure to ambient PM_2.5_, with stronger adverse effects observed in low SES families and in males. Better understanding of the socioeconomic disparities and sexual dimorphism in neurotoxic effects of PM_2.5_ on intellectual development may help elucidate the underlying mechanisms and shed light for targeted and effective interventions.

## Acknowledgements

This study used data from the USC-RFAB twin study. We thank the USC research staff for their assistance in collecting data, and subjects for their participation.

## Supporting Information

**S1 Table.** Descriptive statistics of major demographic characteristics, PM_2.5_ 1-year preceding and IQ scores of three sub-cohorts

**S2 Table.** Population Characteristics at Baseline in Relation to Levels of Verbal IQ

**S3 Table.** Population Characteristics at Baseline in Relation to Levels of Performance IQ

**S4 Table.** Associations between total annual NOx and subscales of IQ

**S1 Fig.** Plot of regression coefficients and 95% confidence intervals for the associations between PM_25_ (1-, 2- and 3-year preceding test) and subscales of IQ from the final-adjusted model

**S1 File. Appendix.** A1. Map of Residential Locations during pre-/early- adolescence and emerging adulthood; A2. Temporal-spatial Modeling of PM_2.5_ Exposure; A3. Relevant Covariates; A4. Summary Table of Air Pollution and IQ Studies

